# Calcium Homeostasis Modulator 2 Constitutes an ATP-regulation Pore in Mitochondria

**DOI:** 10.1101/2024.09.30.615983

**Authors:** Qian-Jin Guo, Tingting Hou, Wen-Jun Xie, Jing-Ruo Zhang, Xin-Lei Ma, Yingna Guo, Xiao-Ting Wang, Li-Peng Wang, Ming-Ao Lu, Zhaofa Wu, Hong-Guang Wang, Yi-Hang Chen, Yu-Long Li, Shi-Qiang Wang

## Abstract

Recent structural analyses showed that the calcium homeostasis modulator-2 (CALHM2) forms a mega channel, but its cellular location and endogenous function are yet unknown. We found that native CALHM2 resides on the mitochondrial inner membrane and constitutes an ATP-regulated ATP release channel. CALHM2 knockdown/knockout decreases cytosolic ATP concentration, and thereby compromises energy-sensitive processes, such as intracellular Ca^2+^ handling. However, CALHM2 loss-of-function elevates ATP concentration in the mitochondrial matrix, dephosphorylates key enzymes in the mammalian target of rapamycin (mTOR) pathway, and promotes longevity in CALHM2 knockout mice. These findings reveal that CALHM2 constitutes a novel regulator of mitochondrial metabolism, which may have important implications in aging and diseases.

## Introduction

The calcium homeostasis modulator (CALHM) family of genes are clustered at 10q24.33 (*CALHM1-3*) and 6q22.1 (*CALHM4-6*) in human genome, and are recent shown to encode mega channels.^1-5^ CALHM1 is predominantly expressed in the cell membrane of neurons, and have been implicated in taste signaling and neurodegenerative diseases.^6-9^ The function of other CALHM proteins are largely unknown.

Recent structural analyses using cryo-electron microscopy have shown that CALHM2 monomers comprise 4 transmembrane domains, the undecamerization of which encompasses a pore of 23-50 Å.^1,2^ Interestingly, a pair of CALHM2 channels with opposite orientation are able to dimerize to form a gap junction-like 22mer.^1,2^ Although overexpression of tagged CALHM2 also creates channels on cell membrane,^1,10^ the cellular location and function of endogenous CALHM2 are poorly understood.

In the present study, we investigated the cellular location and function of endogenous CALHM2, and found that CALHM2 formed metabolism-regulating channels in mitochondria.

## Results

### CALHM2 resides in the mitochondrial inner membrane

CALHM2 is one of the CALHM members in vertebrates, and widely expressed in all tested tissues including heart, liver, spleen, kidney, brain and muscle (Figure 1A). To determine the subcellular location of CALHM2, we stained different types of cells using anti-CALHM2 antibodies. In contrast to the reported localization of overexpressed CALHM2 at the plasma membrane,^1,10^ CALHM2 immuno-signals in COS7 cells exhibited granule patterns that were distributed over the cytosolic space and overlapped with that of mitochondria stained with antibodies against ATP synthase subunit β (ATP5B) (Figure 1B). Similarly, in freshly isolated hepatocytes (Figure 1C), the immunofluorescence of CALHM2 superposed that of the antibodies against ATP5B in mitochondria. These experiments in distinct cell types suggest that CALHM2 is a mitochondrial component.

**Figure 1.**
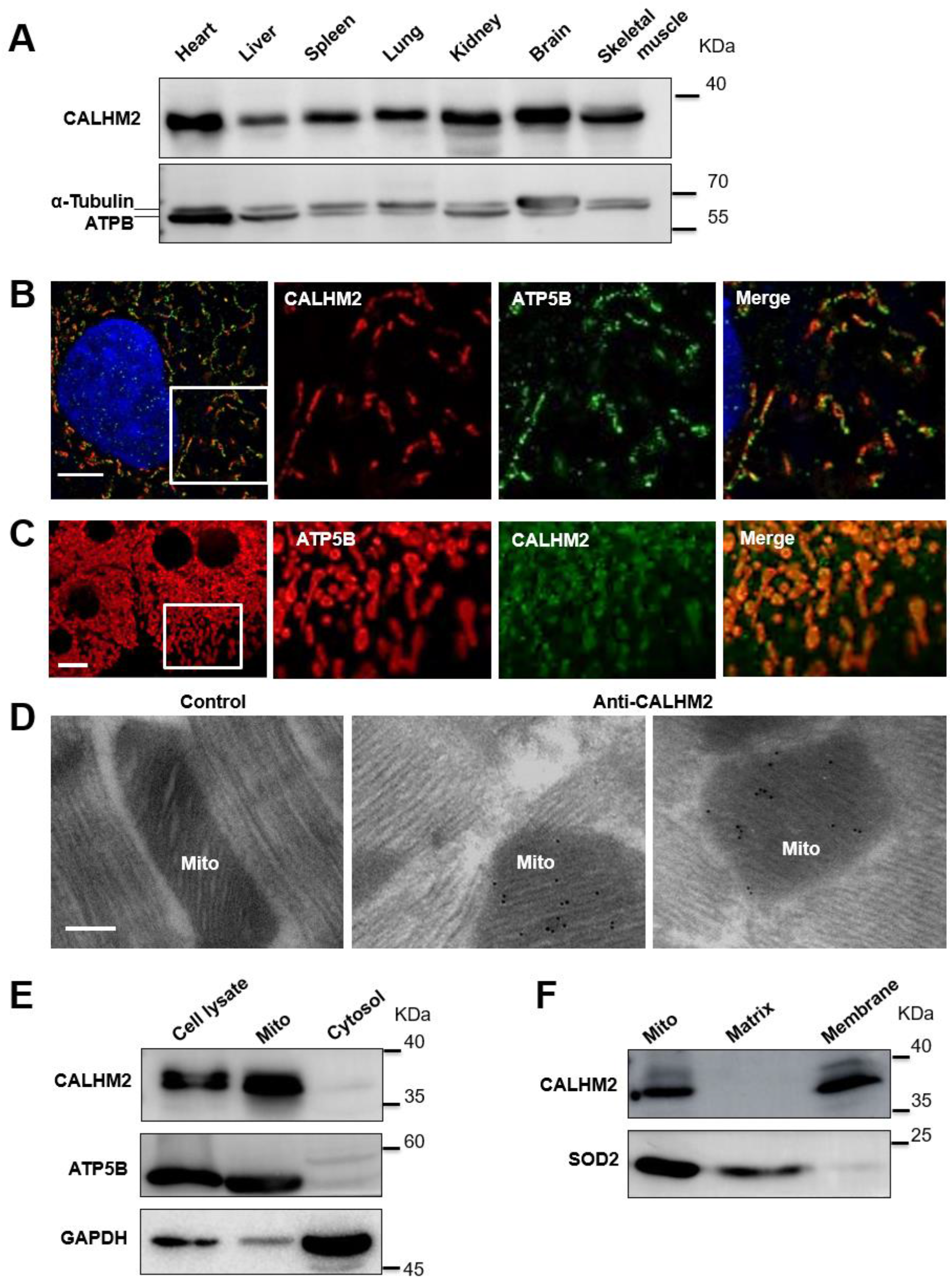
CALHM2 resides in the mitochondrial membrane and regulates cell metabolism. **A**. Typical western blot images showing the expression of CALHM2 in different mouse tissues. α-Tubulin and ATPB were used as controls for cytosolic and mitochondrial proteins, respectively. **B**. Typical super-resolution images acquired with structured illumination microscopy (SIM) showing that the CALHM2 and ATP5B immunofluorescence overlapped with each other in COS7 cells. Scale bar, 5 µm. **C**. Representative confocal images showing colocalization between CALHM2 and ATP5B in mouse hepatocytes. Scale bar, 10 µm. **D**. Immuno-EM images of adult rat cardiomyocytes labeled for CALHM2. Scale bar, 100 nm. **E**. Western blot of CALHM2, ATP5B (a mitochondrial protein) and GAPDH (a cytosol protein) in liver cell lysate and its mitochondrial and cytosolic fractions. **F**. Western blot of CALHM2 and SOD2 (a soluble protein in mitochondria) in mitochondrial lysate and its matrix and membrane fractions. Mitochondria were isolated from liver and subjected to alkali extraction.

In order to determine whether CALHM2 resides on the inner or outer membrane of mitochondria, we performed immuno-electron microscopy (immuno-EM). We found that CALHM2 signals were detected on cristae inside mitochondria (Figure 1D). Considering the transmembrane nature of CALHM2,^1,2^ we inferred that CALHM2 molecules resided in the inner membranes of mitochondria.

We also performed biochemical analysis of cell lysates, which showed that CALHM2 was present in the mitochondria fraction together with ATP5B, but not in the cytosolic fraction (Figure 1E). Furthermore, alkali extraction of purified mitochondria showed that CALHM2 was enriched in the mitochondrial membrane fraction, but not in the matrix fraction (Figure 1F). These results provide further evidence that CALHM2 is a protein in mitochondrial cristae.

### CALHM2 regulates Metabolism

Mitochondrial cristae are crowded with respiratory complexes and metabolic enzymes. To probe the function of CALHM2, we suppressed CALHM2 expression using shRNA in HeLa cells (Figure 2A). Cellular respiration measurement using Seahorse analyzer (Figure 2B) showed that the maximal oxygen consumption rate (OCR, Figure 2C) was significantly decreased. In accordance, luciferase assays showed that the ATP level in CALHM2-knockdown cells was reduced (Figure 2D). These results indicate that CALHM2 plays an important role in the energy metabolism of mitochondria.

**Figure 2.**
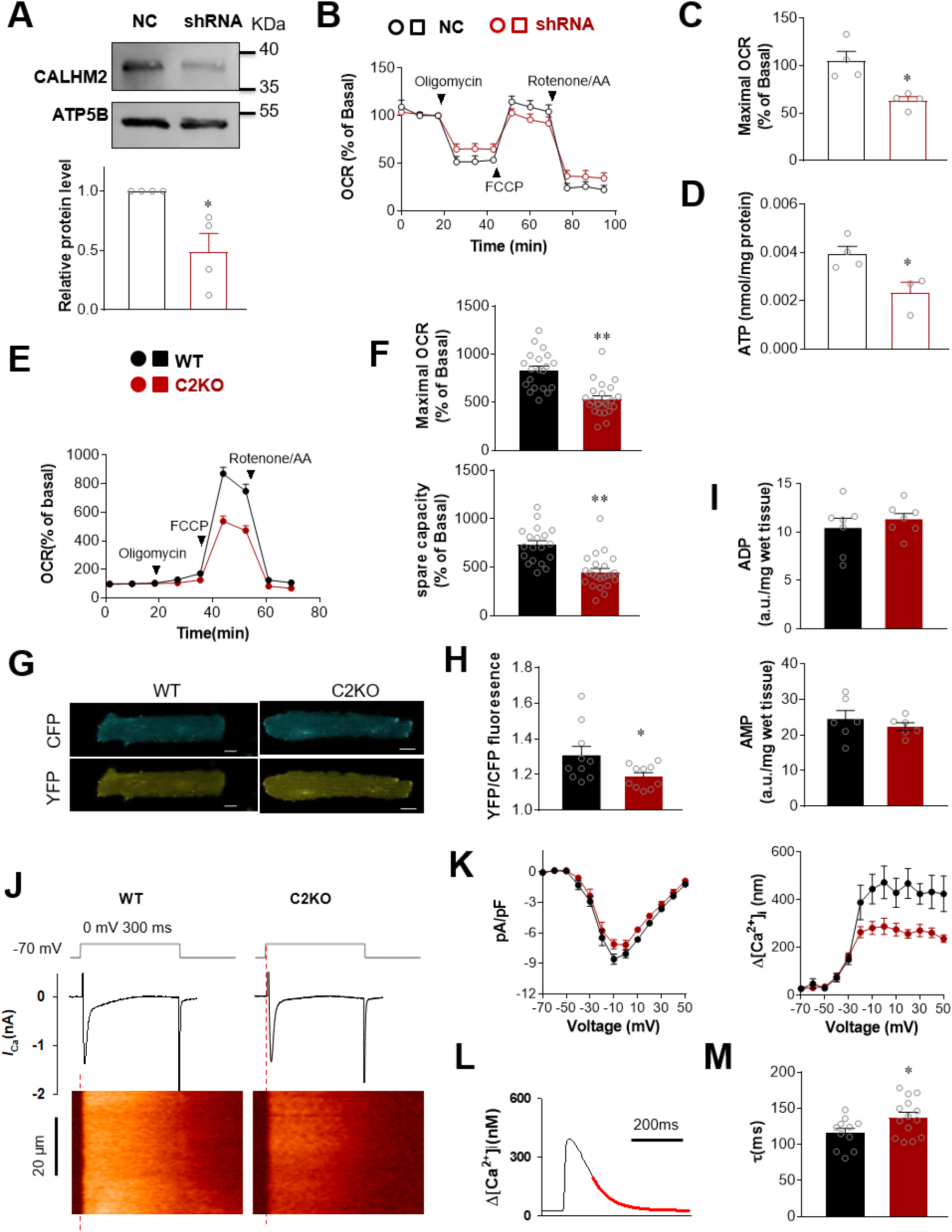
CALHM2 regulates metabolism. **A**. Western blot assay to test CALHM2 knockdown efficiency in HeLa cells of negative control (NC) and shRNA groups. *N* = 4. **B**. Representative respiration traces of NC and CALHM2-knockdown HeLa cells using the XFe24 Seahorse Analyzer. **C**. Comparison of maximal oxygen consumption rate (OCR) between NC and CALHM2-knockdown groups. *N* = 4. **D**. ATP content in NC and CALHM2-knockdown HeLa cells. *N* = 4 and 3 for the NC and shRNA groups, respectively. **E**. Respiration measurement in WT and C2KO cardiomyocytes using an XFe24 Seahorse Analyzer. **F**. Comparison of maximal oxygen consumption rate (OCR) and spare capacity between WT and C2KO cardiomyocytes. **G**. Typical images of ATeam FRET assay in WT and C2KO cardiomyocytes. Scale bar, 10 µm. **H**. Comparison of the YFP/CFP fluorescence ratio between the WT and C2KO groups. *N* = 10 cells from 3 mice in each group. **I**. ADP and AMP content in heart tissues from WT and C2KO mice. *N* =6 and 7, respectively. **J**. Typical recordings of LCC Ca^2+^ currents (*I*_Ca_, upper plots) and Ca^2+^ transients (bottom images) in response to the depolarization from −70 to 0 mV. **K**. *I*_Ca_ density (left) and Ca^2+^ transient amplitude (right) of the WT and C2KO groups. Two-way ANOVA with repeated measures identified a significant difference between WT and C2KO groups. *N =* 9 cells from 3 mice in each group. **L**. Measurement of the rate constant of Ca^2+^ removal. The red curve is the exponential fitting of the decay phase of a Ca^2+^ transient. **M**. The rate constant of Ca^2+^ transient decay of the WT and C2KO groups. \**P* < 0.05. \*\**P* < 0.01.

To confirm genetically that native CALHM2 participates in metabolism regulation, we created a *Calhm2*-knockout (C2KO) mouse model (Extended Data Figure 1). Oxygen consumption measurement in heart cells (Figure 2E) showed that both the maximal OCR and the spare respiratory capacity were decreased in the C2KO group (Figure 2F). These results agreed well with the results in CALHM2-knockdown cell lines, and confirmed that CALHM2 regulates cell metabolism.

We then investigated the role of CALHM2 in ATP regulation. To measure the cytosolic ATP, we loaded cardiomyocytes with ATeam (Figure 2G), a fluorescence resonance energy transfer (FRET)-based indicator for ATP measurement,^11^ which revealed that the cytosolic ATP level, indexed by the YFP/CFP emission ratio, was significantly decreased in isolated C2KO cardiomyocytes (Figure 2H). In contrast with several experimental models in which decreased ATP appears together with increased ADP,^12-14^ we found that the ADP and AMP levels in C2KO cells was unchanged (Figure 2I).

The decrease of ATP supply in C2KO mice would be expected to compromise ATP-dependent processes. Indeed, we found that the Ca^2+^ transients (Figure 2J) triggered by unaltered Ca^2+^ influx through L-type Ca^2+^ channels were decreased significantly in C2KO cardiomyocytes (Figure 2K). In addition, the rate of Ca^2+^ removal, indexed by the rate constant of Ca^2+^ transient decay (Figure 2L), was decreased in C2KO cardiomyocytes in comparison with that of WT cardiomyocytes (Figure 2M). The rate of Ca^2+^ removal reflected the activities of the primary active transport by the sarcoplasmic/endoplasmic reticulum Ca^2+^-ATPase (SERCA) and secondary active transport by the Na^+^-Ca^2+^ exchanger.^15^ The SERCA-mediated Ca^2+^ uptake in turn determines the releasable amount of Ca^2+^ during Ca^2+^ transients.^15^ Therefore, the decreased Ca^2+^ transient amplitude and Ca^2+^ removal rate both reflected that the declined ATP supply in C2KO cells influenced the energy-dependent processes. However, cardiac function was not changed significantly in C2KO mice (Extended Data Figures 1D-E), indicating that the changes in Ca^2+^ handling remained within the stability margin, in which disturbances in microscopic mechanisms are buffered by the system without causing global effects.^16^

### CALHM2 forms an ATP-modulated pore

Surprisingly, in contrast to the decreased ATP level in the cytosol, the ATP level in the mitochondrial matrix was elevated in C2KO cells (Figure 3A). The elevated matrix ATP level and decreased cytosolic ATP level in C2KO cells suggest that CALHM2 is potentially an ATP release channel in mitochondria.

**Figure 3.**
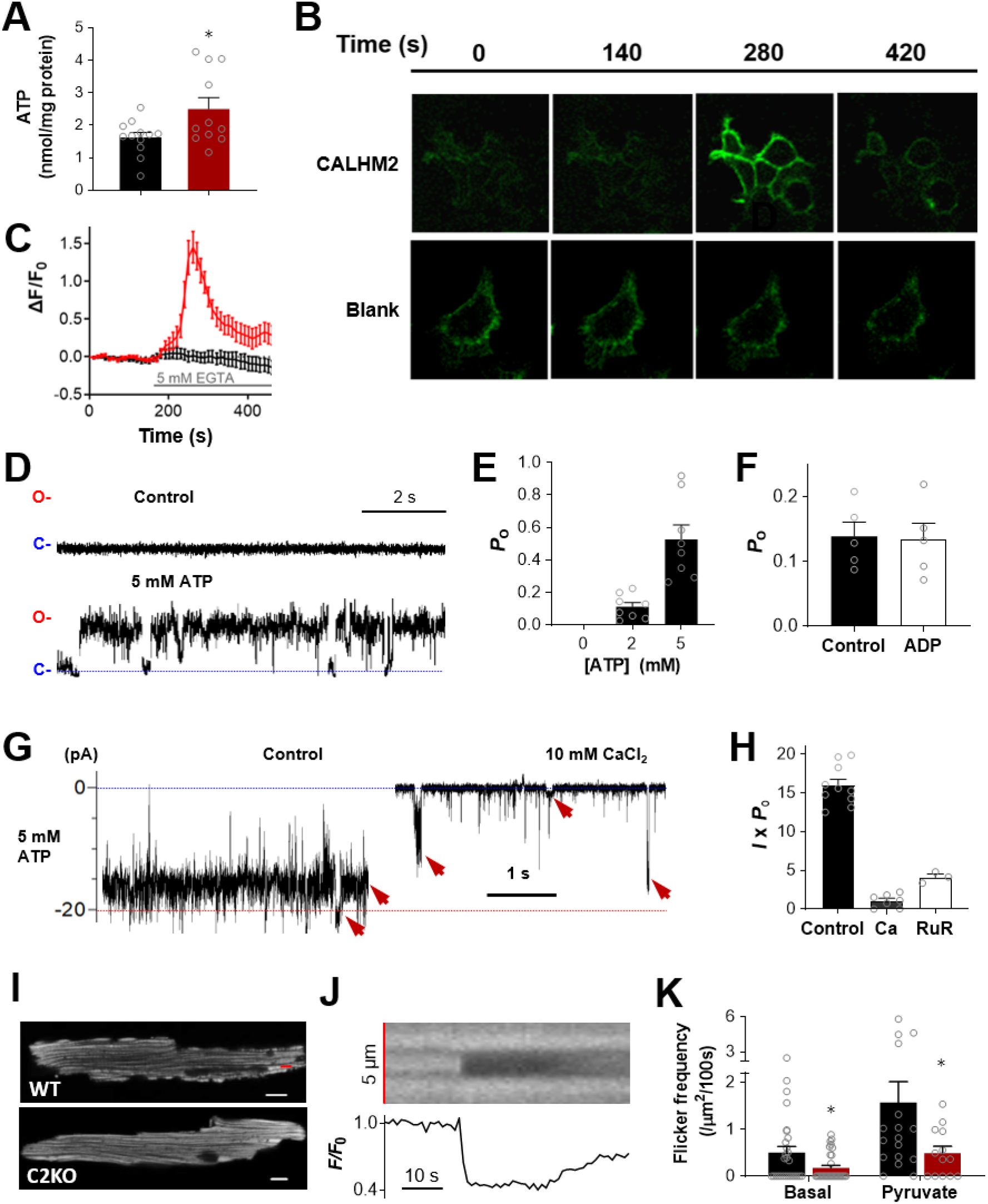
CALHM2 forms an ATP-modulated pore. **A**. ATP content in heart mitochondria from WT and C2KO mice. *N* ≥ 11. **B**. Typical images of GRAB_ATP1.0_ fluorescence in blank cells and cells with CALHM2-flag over-expression before and after the removal of 2 mM Ca^2+^. Scale bar, 20 µm. **C**. Time courses of GRAB_ATP1.0_ fluorescence trace before and after 2 mM Ca^2+^ removal in blank cells and cells with CALHM2-flag over-expression. **D**. Representative recording of CALHM2 channels in 0 mM (control) and 5 mM ATP environments. **E**. Comparison of the open probability (*P*_O_) of CALHM2 channels in 0 mM, 2 mM and 5 mM ATP environments. *N* = 8. **F**. Comparison of the *P*_O_ in the presence of 2 mM ATP and 2 mM ATP + 1 mM ADP. *N* = 5. **G**. Typical recording of CALHM2 mega channel activity in 5 mM ATP before and after the addition of 10 mM CaCl_2_. Red arrows denote different levels of channel currents. **H**. Conductance of CALHM2 channels in the control, 10 mM CaCl_2_ and 100 µM RuR groups. *N* = 10, 7 and 3 in the control, Ca and RuR groups, respectively. **I**. Typical confocal images of TMRM fluorescence showing the ΔΨ_m_ in WT and C2KO cardiomyocytes. **J**. The time-lapse images and time course of ΔΨ_m_ at the red line area in **I**, showing a typical ΔΨ_m_ flicker. **K**. Comparison of the frequency of ΔΨ_m_ flickers in WT and C2KO cardiomyocytes with and without pyruvate treatment. *N* = 13 to 31 cells from 3 mice in each group. \**P* < 0.05.

Recently reported structural data^1-5^ show that the pore of a CALHM2/CALHM6 undecamer is wider than that of the ATP-permeable CALHM1 octamer. To confirm that CALHM channels are permeable to ATP, we expressed flag-tagged CALHM2 on the plasma membrane of HeLa cells as reported previously, and imaged the ATP release activity using a newly-developed extracellular ATP sensor, GRAB_ATP1.0_.^17^ It is recently reported that CALHM2 channels expressed on cell membrane is activated when extracellular Ca^2+^ is removed.^1^ We found that the GRAB_ATP1.0_ fluorescence (Figure 3B) was transiently increased in the CALHM2-flag group, but not in the control group, when the extracellular Ca^2+^ was switched from 2 to 0 mM (Figures 3C). As GRAB_ATP1.0_ only detects extracellular ATP,^17^ these findings indicate that ATP is released from the cytosol via overexpressed CALHM2 channels.

In order to understand how CALHM2 channels are regulated, we incorporated CALHM2 molecules into lipid bilayers and studied the effects of mitochondrial metabolites on the electrophysiological properties of CALHM2 channels. In the presence of 135 mM K^+^ and 15 mM Na^+^ with 2 mM ATP, we successfully detected channel gating activity (Figure 3D) with a unitary conductance of ∼33 pS. When the ATP concentration was increased to 5 mM, the open probability of CALHM2 channels was increased significantly (Figures 3E). In contrast to its ATP sensitivity, the channel open probability in the presence of 2 mM ATP was not changed by additional 1 mM ADP (Figure 3F). These results indicate that the activity of CALHM2 channels is regulated by ATP.

In the presence of 5 mM ATP, we often recorded 18-22 pA currents at a bias of ±50 mV (Figure 3G). This mega conductance was one-order-of-magnitude larger than the unitary conductance and was rarely observed in 2 mM ATP. Its 360-440 pS conductance was comparable to that of pannexin 1 and P2X_7_ receptors, which are ATP-permeable channels.^18-21^

Agreeing well with recent reports^1^, both Ca^2+^ (10 mM, Figure 3G right) and ruthenium red (RuR, 100 µM) suppressed CALHM2 channel conductance (Figure 3H). In most recordings of the mega conductance, we observed different levels of channel openings (e.g., red arrows in Figure 3G). The variation of the mega conductance should reflect the asynchronous activation of individual protomers in a CALHM2 channel.

To probe the *in situ* activity of CALHM2 channels, we utilized tetramethylrhodamine methyl ester (TMRM, Figure 3I) to monitor the mitochondrial membrane potential (ΔΨ_m_). We inferred that if the CALHM2 channel exists in mitochondria then its opening with mega conductance would transiently decrease ΔΨ_m_. In WT cardiomyocytes, the transient decrease of ΔΨ_m_ in individual mitochondria (Figure 3J), also known as mitochondrial flickers,^22-24^ occurred stochastically. In contrast, the ΔΨ_m_ flickers were suppressed by 65% after knockout of CALHM2 (Figure 3K left bars). Pyruvate is an immediate metabolite that feeds the tricarboxylic cycle and boost mitochondrial ATP production.^25,26^ While pyruvate increased ΔΨ_m_ flickers in both WT and C2KO groups, the pyruvate-elicited flicker frequency was decreased by 77% in C2KO cardiomyocytes (Figure 3K right bars). These results indicated that approximately 2/3 of mitochondrial flicker events were attributable to the opening of CALHM2 channels *in situ*.

### CALHM2 modulates the activity of metabolic proteins

Blue native polyacrylamide gel electrophoresis (BN-PAGE, Figure 4A) showed that C2KO did not influence the apparent composition of respiratory complexes (Figure 4B). However, the enzymatic activity of Complex V, but not that of Complexes I and II, was significantly decreased in the C2KO group (Figures 4C-E), agreeing well with decreased OCR shown above (Figures 2B-G).

**Figure 4.**
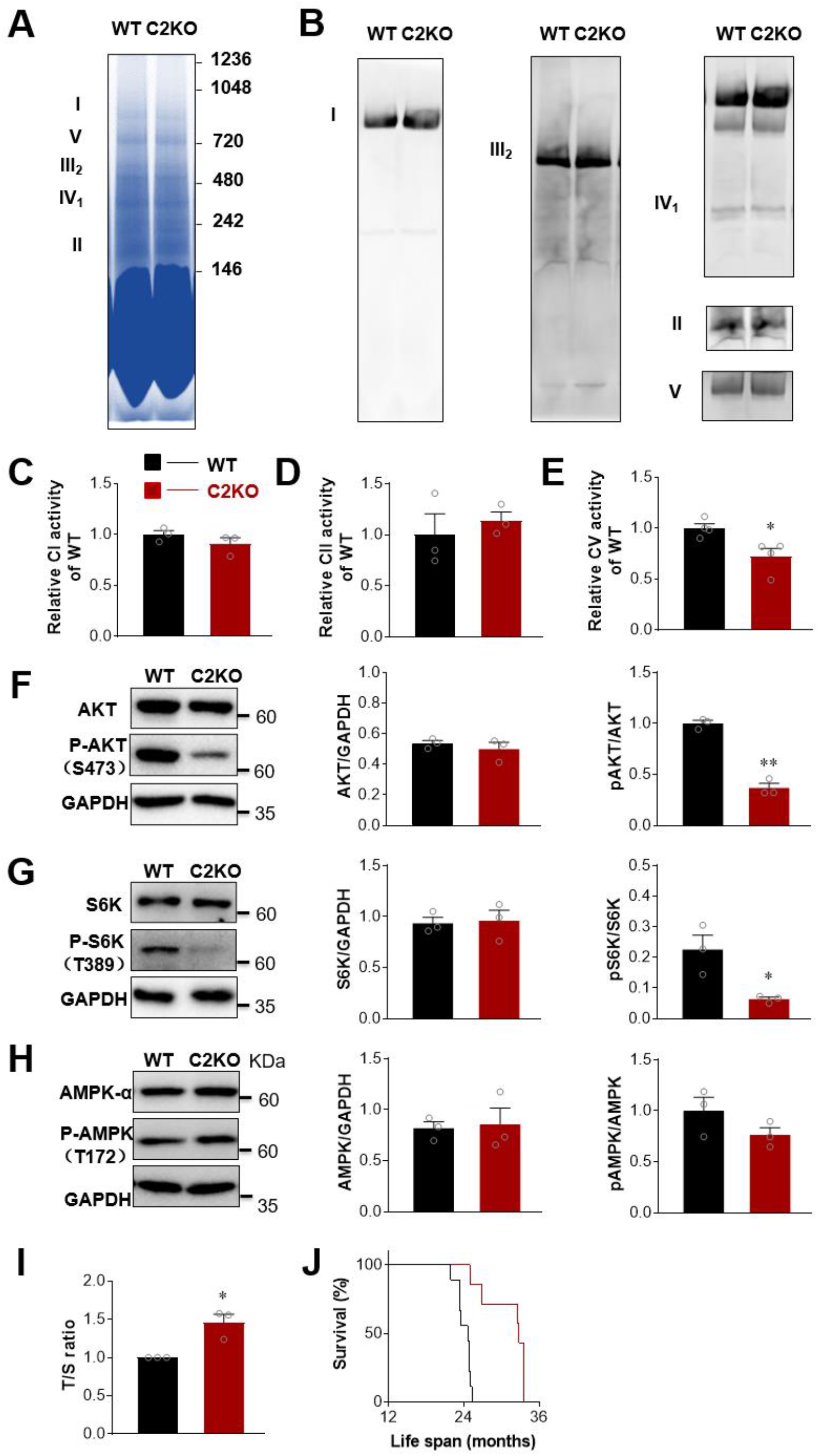
CALHM2 modulates mTOR signaling pathway and impacts on mouse lifespan. **A**. Typical blue native gel of Complexes I, II, III, IV and V. **B**. Western blot images for mitochondrial Complex I (anti-NDUFB8), II (anti-SDHA), III2 (anti-UQCRC1), IV (anti-COX4) and V (anti-ATP5B). **C-E**. Comparison of mitochondrial Complex I, II and V activity between wild-type (WT) and C2KO groups. *N* = 4 in each group. **F**. Typical Western blot images (left) and the quantification of AKT (middle) and phosphorylated AKT (p-AKT, recognized by an antibody against S473, right) in WT and C2KO mouse liver tissues, normalized by GAPDH. *N* = 3. **G**. Typical Western blot images (left) and quantification of S6K (middle) and phosphorylated S6K (p-S6K, recognized by an antibody against T389, right) in WT and C2KO mouse liver tissues, normalized by GAPDH. *N* = 3. **H**. Typical Western blot images (left) and the quantification of AMPK-α (middle) and phosphorylated AMPK-α (p-AMPK, recognized by an antibody against T172, right) in WT and C2KO mouse liver tissues, normalized by GAPDH. *N* = 3. **I**. Measurement of the ratio between the amount of the telomere repeat amplification product to that of 36B4 (RPLP0) (T/S ratio) in WT and C2KO mouse liver. *N* = 3. **J**. Kaplan–Meier survival analysis of WT and C2KO mice observed over three years. *N* = 13 female WT and 11 female C2KO mice. \**P* < 0.05. \*\**P* < 0.01.

Metabolism suppression is expected to influence the status of key proteins in metabolic pathways. We found that the phosphorylation of Akt, also known as protein kinase B,^27^ was suppressed by 66% in the C2KO group (Figure 4F). Akt regulates the target of rapamycin (TOR, or its mammalian version mTOR) signaling and diverse cellular functions.^27-30^ The decrease of Akt phosphorylation at S473, which links the signaling cascades between mTOC Complexes 2 (mTORC2) and 1 (mTORC1),^26,30^ suggesting that mTOR signaling was suppressed in the absence of CALHM2. Indeed, the ribosomal S6 kinase (S6K), a well-characterized mTOR downstream target,^31^ were dephosphorylated (Figure 4G). Agreeing with the unchanged AMP level (Figure 2I), the phosphorylation of the AMP-activated protein kinase (AMPK) was not changed in the C2KO group (Figure 4H). These results indicated that the mTOR signaling pathway, but not that of AMPK, was strongly inhibited in C2KO cells.

It is found in *Caenorhabditis elegans* that the TOR activity and aging process are tightly related to mitochondrial respiration and ATP production.^32-35^ With the down-regulated mTOR signaling in C2KO mice, we found that the ratio between the amount of the telomere repeat amplification product to that of 36B4 (RPLP0) (T/S ratio), which reflects the average telomere length,^36,37^ was higher in 13-month old C2KO mice than in WT mice (Figure 4I), suggesting that the aging process was delayed in C2KO cells. In accordance, three years of observation showed that the lifespan of C2KO was extended by 26% (Figure 4J). This finding suggests that the “ATP↓-TOR↓-longevity” principle revealed in worm models^33^ also applies in mammals.

## Discussion

In the present study, we characterized CALHM2 as an ATP-modulated ATP-permeable channel in the mitochondrial inner membrane. Previous reports have shown that CALHM2 labeled with flag^10^ or green fluorescent protein (GFP)^1^ was expressed on the cell membrane, and that GFP-linked CALHM2 cannot form gap junction-like channels.^1^ We also observed that flag-tagged CALHM2 was partially expressed on cell membrane. These phenomena suggested that the tags may interfere with the location of CALHM2 in mitochondria. In the present study, we provide several lines of evidence showing that native CALHM2 is a mitochondrial protein: 1) The intracellular distribution of CALHM2 overlapped that of mitochondria in different cell types; 2) immuno-EM imaging revealed CALHM2 inside mitochondria; 3) biochemical analyses showed that CALHM2 exists in the mitochondrial membrane, but not in the cytosol or mitochondrial matrix; 4) knockdown and knockout of CALHM2 influenced the oxygen consumption; 5) the ΔΨ_m_ flickering of mitochondria was largely suppressed in C2KO cardiomyocytes; and 6) ATP concentration was decreased in cytosol but increased in mitochondria. These facts provide firm evidence that CALHM2 forms an ATP-release channel in mitochondria. The role of CALHM2 in ATP supply also explains the reported decrease of ATP release in CALHM2-knockdown/knockout astrocytes as well as its neural behavioral consequences.^10^

Mitochondria are known to release ATP mainly through the adenine nucleotide transporter (ANT).^38-40^ In addition to the ANT, the mPTP act as a mega channel in mitochondrial inner membrane, which is also permeable to ATP.^41-45^ Given its ATP-sensitivity, the CALHM2 channel may act as a “pressure-relief valve” that releases ATP in a burst manner when the matrix ATP level reaches a certain threshold, which differs from the relatively smooth, house-holding ATP release by ANT. Therefore, we propose that CALHM2 constitutes a third mechanism for ATP release from mitochondria. Given the existence of different ATP release mechanisms, although CALHM2 knockout moderately affected ATP-dependent processes, such as intracellular Ca^2+^ handling, major biological processes, including heart function, remain in a healthy range.

With the decrease of cytosolic ATP in C2KO mice, the phosphorylation of key proteins in the mTOR pathway was suppressed, and the lifespan of C2KO mice was significantly longer than WT mice. These findings agree well with the worm experiments that α-ketoglutarate treatment decreases cytosolic ATP, suppresses TOR signaling and leads to longevity in *C. elegans*.^33^ Our findings of the longevity in C2KO mice linked the reduction of cytosolic ATP to mTOR suppression and lifespan extension in mammals, which may provide a possible explanation that calorie restriction suppresses the mTOR pathway and extends the lifespan in mammals.^46-47^ AMPK is another conserved sensor of cellular energy status linked to longevity.^47,48^ In our mice model, C2KO decreased cytosolic ATP without altering ADP and AMP levels. As AMPK is an AMP-activated protein kinase,^47,48^ it is not surprising that the AMPK phosphorylation activity was not changed in C2KO mice. These results provide direct evidence that the TOR-dependent but AMPK-independent longevity found in worms^33^ is also inducible in mammals.

In summary, we found that CALHM2 forms ATP-modulated ATP permeable channels on the mitochondrial inner membrane that modulate cell metabolism. Cells from CALHM2 knockout mice exhibited decreased ATP in the cytosol but increased ATP in mitochondrial matrix, which was attributable to decreased mitochondrial ATP release. The decrease in cytosolic ATP suppressed mTOR signaling and extended the lifespan in C2KO mice. These results reveal that CALHM2 is a novel tuner of mitochondrial metabolism, which may have important implications in the understanding of aging and metabolic diseases.

## Methods

### Animal generation

*Calhm2*^*+/-*^ mice were created in the Model Animal Research Center of Nanjing University. Mice were generated in the C57BL/6 mice background. The animal handling procedures (No. LSC-Wangsq-1) were approved by the Institutional Animal Care and Use Committee (IACUC) at Peking University accredited by AAALAC international. The animal experiments conform to the Guide for Care and Use of Laboratory animals published by the US National Institutes of Health (NIH Publication No.85-23, revised 2011). All mice were housed in a pathogen-free barrier environment, and were kept on a 12-h light–dark cycle at 21-23°C.

### Mitochondria isolation and alkali extraction analysis

A mitochondria isolation kit (Abcam) was used to perform mitochondrial isolation. Briefly, fresh tissues and cells were washed twice with 1.5 ml of wash buffer and were homogenized in isolation buffer using a pre-chilled Dounce homogenizer. The homogenate was centrifuged at 1,000 × *g* for 10 minutes at 4 °C. Next, the supernatant was centrifuged at 12,000 × *g* for 15 minutes at 4 °C, after which the pellet (mitochondria) and supernatant (cytosol) were collected.

For the alkali extraction analysis, the isolated mitochondria were re-suspended in alkali extraction reagent (0.1 M Na_2_CO_3_, pH 10.5) and incubated for 1 h on ice. The suspension was then centrifuged at 100,000 × *g* for 30 min at 4 °C. The supernatant was collected as the mitochondrial soluble fraction, and the pellet was collected as the mitochondrial membrane fraction.

### Western blot analysis

Tissues were homogenized in RIPA buffer (Beyotime) containing phenylmethylsulfonyl fluoride (PMSF, Thermo Fisher). The lysates were agitated and centrifuged at 4 °C. The supernatant protein mixture was separated by 12% SDS-PAGE analysis and subjected to western blot analysis with antibodies against anti CALHM2 (Abcam,ab121446), ATP5B (Abcam, ab14730), β-actin (Cell Signaling, 4970), AMPKα (Cell Signaling, 5831T), P-AMPKα T172 (Cell Signaling, 2535T), AKT (Cell Signaling, 4691S), P-AKT S473 (Cell Signaling, 4060S), S6K (Cell Signaling, 9202S), P-S6K T389 (Cell Signaling, 9234), SOD2 (Abcam, ab13533), α-Tubulin (Sigma, T9026), ATP5A1(Abcam, ab110273), NDUFB8 (Abcam, ab110242), UQCRC1 (Abcam, ab110252), COXIV (ProteinTech, 11242-1-AP) and SDHA (Abcam, ab14715). Horseradish peroxidase-conjugated goat anti-rabbit or anti-mouse antibody (ThermoFisher, 1:5000) were used for detection. The horseradish peroxidase-conjugated GAPDH antibody (Shanghai Kangchen) was used to detect GAPDH content as the loading control.

### Immunofluorescence staining

Cells were fixed in 4% paraformaldehyde for 15 min and then washed in phosphate-buffered saline (PBS). The cells were permeabilized by 0.5% Triton-X for 15 min, incubated with 0.5% Triton-X and 10% goat serum albumin for 30 min, washed, and incubated with primary antibodies against CALHM2 (Abcam), ATP5B (Abcam) or SDHA (Abcam) overnight at 4 °C. Next, the cells were washed in PBS and incubated with different secondary antibody for 2 h at room temperature. Confocal images were acquired using a Zeiss microscope (LSM 710, Zeiss, Jena, Germany).

### Immuno-electron microscopy

Cardiomyocytes grown on glass coverslips were fixed in 3% paraformaldehyde and 0.1% glutaraldehyde in 0.1 M sodium cacodylate (pH 7.2-7.4) for 4 h at 4 °C and washed with 0.1 M sodium cacodylate buffer containing 4% sucrose and 0.1 M glycine (pH 7.4) at 4°C. After dehydration in a graded ethanol series, the cells were embedded in LR White (London Resin Co). Ultrathin sections were attached to square nickel mesh grids and blocked for 5 min with 1% bovine serum albumin (Sigma) in PBS. The sections were incubated for 2 h with anti-CALHM2 antibody diluted 1:50 with PBS. After washing with PBS, the sections were incubated for 1 h with donkey anti-rabbit IgG coupled to 10-nm gold particles (Jackson ImmunoResearch) diluted 1:50 with PBS. The sections were contrasted with uranyl acetate and lead citrate, after which they were observed under an FEI Tecnai G^2^ 20 Twin system electron microscope. In the control group, normal mouse serum was substituted for the primary antibody.

### Mitochondrial respirometry

Respirometry of adult cardiomyocytes was performed using an XFe24 Extracellular Flux Analyzer (Seahorse Biosciences). Freshly-isolated cardiomyocytes were seeded at 4000 cells/well in 24-well XFe microplates coated with 10 μg/ml laminin. After 1 h incubation in Tyrode solution in a 5% CO_2_ incubator at 37 °C, the medium was replaced with XF base medium supplemented with 4 mM glucose and 1 mM pyruvate, after which the cells were incubated in a CO_2_-free incubator at 37 °C for another 1 h to allow temperature and pH equilibration. The baseline O_2_ consumption rate (OCR) was measured, after which sequential injections of oligomycin (1 μM) were performed to measure the ATP-linked OCR, the oxidative phosphorylation uncoupler carbonyl cyanide 4-(trifluoromethoxy) phenylhydrazone (FCCP) (0.2 μM) to determine maximal respiration, and rotenone (1 μM) and antimycin A (1 μM) to determine the non-mitochondrial respiration. Experimental treatments were performed on 4-11 wells of each plate as technical replicates, and each experiment had at least 3 replicates.

### Measurement of mitochondrial membrane potential ΔΨ_m_

TMRM (ThermoFisher) stock solution was prepared as described in the manual (Catalog number T668). The stock solution was diluted to a final concentration of 100 nM. Cells were incubated with the TMRM stock solution for 10 minutes at room temperature. For ΔΨ_m_ detection, time-lapse two-dimensional (x × y) images were captured and 100 frames of 512 × 512 (x × y) pixels were collected at a rate of 1 s/frame in bidirectional scanning mode in a given region. The images were acquired by excitation at 543 nm, and emissions were measured at >560 nm.

### Planar lipid bilayer analysis

Purified CALHM2 protein was used for the single channel recording. Proteins were fused into planar lipid bilayers formed by painting a lipid mixture of phosphatidylethanolamine (DOPE) and phosphatidylcholine (DOPC) (Avanti Polar Lipids) at a 3:1 ratio in decane across a 200-µm hole in a polystyrene partition separating the internal and external solutions in poly-sulfonate cups (Warner Instruments). The cis-chamber (1.0 ml), representing the luminal compartment, was connected to the head stage input of a bilayer voltage clamp amplifier. The trans-chamber (1.0 ml), representing the cytoplasmic compartment, was held at virtual ground. The Solution in the trans-chamber was 135 mM KCl, 15 NaCl, and the solution in cis-chamber was 10 mM HEPES (pH 7.4). Currents were recorded every minute after application of the voltage to the cis-side. Single-channel currents were recorded using a BC-535 Bilayer Clamp Amplifier (Warner Instruments, LLC, CT), filtered at 1 kHz using a Low-Pass 8-pole Bessel Filter (Warner Instruments, LLC, CT), and digitized at 4 kHz. All experiments were performed at room temperature (23 ± 2 °C). The pH of the solution was adjusted with choline. The single-channel Po was determined over 1 minute of continuous recording using the 50% threshold analysis method. The recordings were analyzed using Clampfit 10.1 software (Molecular Devices).

### Whole-cell patch clamp

Cardiomyocytes were bathed in an external recording solution containing (in mM) 137 NaCl, 4 KCl, 2 CaCl_2_, 1 MgCl_2_, 1.2 NaH_2_PO_4_, 10 glucose, 0.02 tetrodotoxin, and 10 HEPES, pH 7.35 adjusted with NaOH. The internal pipette solution contained (in mM) 110 CsCl, 6 MgCl_2_, 5 Na_2_ATP, 15 TEA-Cl, 10 HEPES, and 0.2 fluo-4 pentapotassium, pH 7.2 adjusted with CsOH. The Ca^2+^ current (*I*_Ca_) was activated by 300 ms pulses from -70 mV to 0 mV at 10-s intervals and recorded using an EPC7 amplifier (List Medical Electronic, Germany). Ca^2+^ transients were recorded by confocal line scanning using a Zeiss LSM-510. Experiments were performed at room temperature (∼25 °C).

### BN-PAGE analysis of mitochondrial supercomplexes and complexes

BN-PAGE was conducted using the NativePAGE^TM^ system (Invitrogen). Briefly, isolated cardiac mitochondria were solubilized by digitonin (4 g/g protein) for 15 min on ice. After centrifugation at 15,000 rpm at 4 °C for 30 min, the supernatants were collected and the protein concentration was determined by BCA analysis (ThermoFisher). Coomassie blue G-250 (Invitrogen) was added to the supernatant to obtain a dye/detergent mass ratio of 4/1, and the protein was loaded into a 4-16% non-denaturing polyacrylamide gel (Invitrogen). After electrophoresis, proteins were transferred to a PVDF membrane and probed with specific antibodies against subunits of Complexes I (NDUFB8), II (SDHA), III (UQCRC1), IV (COX IV), and V (ATP5B or ATP5A). Blots were visualized using secondary antibodies conjugated with IRDye (LI-COR, Lincoln, NE, USA) and an Odyssey imaging system (LI-COR).

### ATP, ADP and AMP measurement

Heart tissue was excised and lysed with Tris-phenol, after which the mitochondrial ATP was extracted with cold 2.5% trichloroacetic acid. The ATP content was measured by luciferase assay according to manufacturer’s instructions (Promega, Madison, WI, USA). ADP and AMP contents were measured by colorimetric assay according to the manufacturers’ instructions (Abcam and Biovision, respectively). The cytosolic ATP was measured by expressing ATeam in isolated cardiomyocytes. Briefly, the ATeam fluorescence was acquired by excitation at 405 nm and emission at 450-515 nm (for CFP) and 515-569 nm (for YFP), respectively. The ATP release was measured by expressing GRAB_ATP1.0_ in HeLa cells. The GRAB_ATP1.0_ fluorescence was recorded with excitation at 488 nm and emission collection at 506-702 nm.

### Measurement of average telomere length

Liver tissue was used to extract the DNA following the phenol-chloroform extraction method and genome DNA was used for real-time PCR amplification with Brilliant II SYBR Green QPCR master mix (Stratagene) on the Mx3000p Real-Time PCR System (Stratagene).The primers for measure the telomere repeat were (forward: 5’-CGGTTTGTTTGGGTTTGGGTTTGGGTTTGGGTTTGGGTT; reverse: 5’-GGCTTGCCTTACCCTTACCCTTACCCTTACCCTTACCCT); The primers for the single copy gene-36B4 were (forward: 5’-ACTGGTCTAGGACCCGAGAAG; reverse: 5’-TCAATGGTGCCTCTGGAGATT).

### Statistical analysis

Results are expressed as mean ±SE. Statistical analysis was performed using Student’s t-test for normal test-past data, the Mann–Whitney rank sum test, and two-way ANOVA with repeated measures or the log-rank test for survival analysis of lifespan. A value of *P* <0.05 was considered significant.

## Acknowledgment

We thank Drs. Ying-Chun Hu, Chun-Yan Shan and Xue-Mei Hao in the National Center for Protein Sciences at Peking University for assistance with the electron microscopy and optical imaging. This study was supported by the National Natural Science Foundation of China (92254301, 32250003, 32200930 and 32230048).

---

**Extended Data Figure 1.**
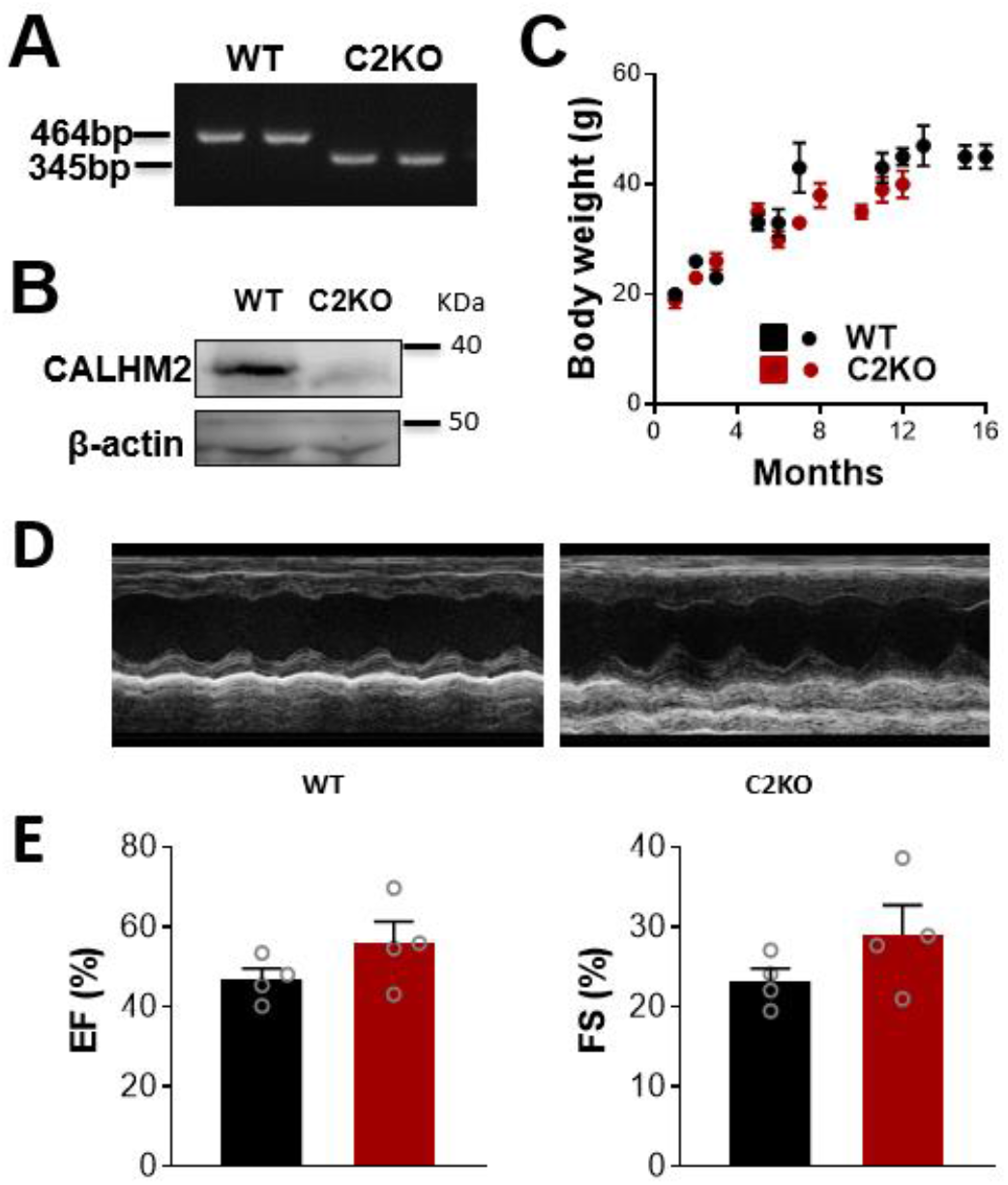
Characterization of the C2KO mice. **A**. Typical genotyping images for WT and C2KO mice. The band size for the WT and C2KO groups was 464 bp and 345 bp, respectively. **B**. Typical Western blot images for CALHM2 expression in WT and C2KO groups. **C**. Developmental changes in the body weight of WT and C2KO mice. **D**. Typical echocardiography images of WT and C2KO mouse hearts. **E**. Comparison of the ejection fraction (EF, left) and fractional shortening (FS, right) between WT and C2KO mouse hearts. *N* = 4.

